# Viral Sequence Identification in Metagenomes using Natural Language Processing Techniques

**DOI:** 10.1101/2020.01.10.892158

**Authors:** Aly O. Abdelkareem, Mahmoud I. Khalil, Ali H. A. Elbehery, Hazem M. Abbas

## Abstract

Viral reads identification is one of the important steps in metagenomic data analysis. It shows up the diversity of the microbial communities and the functional characteristics of microorganisms. There are various tools that can identify viral reads in mixed metagenomic data using similarity and statistical tools. However, the lack of available genome diversity is a serious limitation to the existing techniques. In this work, we applied natural language processing approaches for document classification in analyzing metagenomic sequences. Text featurization is presented by treating DNA similar to natural language. These techniques reveal the importance of using the text feature extraction pipeline in sequence identification by transforming DNA base pairs into a set of characters with a term frequency and inverse document frequency techniques. Various machine learning classification algorithms are applied to viral identification tasks such as logistic regression and multi-layer perceptron. Moreover, we compared classical machine learning algorithms with VirFinder and VirNet, our deep attention model for viral reads identification on generated fragments of viruses and bacteria for benchmarking viral reads identification tools. Then, as a verification of our tool, It was applied to a simulated microbiome and virome data for tool verification and real metagenomic data of Roche 454 and Illumina for a case study.

## 1. Introduction

On our planet, there are more than 1.2 million species of organisms documented in databases. In spite of that, these species are only around 10% of the actual number of species on Earth and in the ocean Mora et al. (2011). Microorganisms are microscopic organisms including all unicellular and some multicellular organisms, Prokaryotes are unicellular organisms such as bacteria and archaea. There are six types of microorganisms which are viruses, protozoa, bacteria, archaea, algae, and fungi. They are found everywhere on Earth and the ocean and they are critical in our survival.

The prokaryotic cell is 0.5 micrometer in diameter or less and it contains DNA floating and not bound by a nuclear membrane. Their chromosomes contain histone-like proteins and they reproduce by binary fission. They also do not contain mitochondria and lysosymes. On the other hand, the eukaryotic cell size is more than 5 micrometer. It also contains the Nucleus where the DNA is bounded by its membrane. Their chromosomes contain histone proteins and They reproduce by mitotic and meiotic division. The cells have mitochondria and lysozymes as well. This study focuses on prokaryotic microorganisms (e.g. bacteria and archaea) and viruses.

The majority of prokaryotic organisms are bacteria and archaea are the smallest independently living single-celled organisms. Bacteria are unicellular and microscopic organisms that reproduce by binary fission. They are found everywhere, in water, soil, human gut, and animal skins. The number of estimated bacteria on Earth is 5 quintillion (10^30^) Whitman et al. (1998). Bacterial cells length are between 0.3 and 5.0 micrometers and most of them have either spherical shaped (cocci) or rod-shaped (bacilli). Their genomes contain from 200 to a few thousand genes. Another type of microorganisms is Archea, which exists alongside bacteria and it is very difficult to differentiate between bacteria and archaea based on their structure and appearance.

Viruses consist of genetic materials surrounded by a coat of proteins, called capsid and some viruses have their genetic material inside a membranous envelope. Viruses can only replicate inside living host cells. The tiniest viral genomes have only three genes and its length is 20 nanometers in diameter. While the largest have up to 2000 genes and its diameter is 1500 nanometers which are very tiny to be visible under the light microscope. Viruses are classified according to their genetic materials which might be single or double-stranded DNA or RNA. Their genome might be a single linear or circular molecule. It classified according to their capsid too as it may be rod-shaped (helical viruses) or polyhedral (Icosahedral viruses). They lack metabolic enzymes and translational machinery such as ribosomes for making proteins.

Viruses that infect bacteria called phages and they are abundant in various microbiome communities. The viral infection starts when a virus binds to a host cell and its genome integrates with the host cell genome. The integrated viral DNA is called a provirus. So, it is very reasonable to think that isolated viruses are just a package of genes moving from one host cell to another.

Viruses have an essential role in various microbial communities, and virus-host interaction can change many ecosystems such as human health and aquatic life. In marine ecology, it is estimated that that one drop of water contains about 10 million viruses. Phages are essential to the marine community because they infect and destroy some bacterial communities, which have an important role in recycling carbon and stimulate algal growth by the released organic molecules from bacteria. They also destroy harmful bacteria which kill other marine organisms Rohwer et al. (2009).

Viral diversity and viral-host interactions in microbial communities are studied using Isolation and culturing techniques. Unlike other organisms, viruses need a host cell for replication. They can culture the infected host cells and then use isolation techniques to study viruses. Viruses particles can be isolated by either filtration or centrifugation. Simply, any particle has a size less than the filter pore size can be collected. In order to collect viruses, we need the filter pore size to be from 20 nanometers to 1500 nanometer.

Culturing techniques encounter some limitations due to absence of universal marker gene for viruses. It is reported that the available viruses genomes in public databases(e.g NCBI RefSeq) is approximately 5% of known species of prokaryotic organisms Roux et al. (2015b). Furthermore, the difficulty of pure culture isolation lead to a small population of microbial organisms that were identified. So, this technique is limited to less than 1% of host cells and is biased to specific organisms as well. Labonté et al. (2015).

Metagenomics is a so is an analysis of the genetic information of the collective genomes of the microbes within a given environment based on its sampling regardless of cultivability of the cellsIzard and Rivera (2014). Culture the majority of microorganisms in the environment is not available through standard techniques due to the difficulty of culturing microorganisms,. However, uncultured organisms are diverse and it is very important to study their diversity and the relation between them and the cultured one Riesenfeld et al. (2004). Therefore, the culture-independent genomic analysis demonstrates a promising understanding of different microorganisms as help researchers to know the identity of microorganisms in the collection and their potential functional characterization. Sequencing viruses with a collection of other organisms is a challenging problem Edwards and Rohwer (2005). Some viruses are not associated with their host genomes such as free viruses in the environment and viruses that kill their host cells. Contaminated prokaryotic population with viral sequences is challenging to many scientists. It is reported that 4-17% of human gut metagenomes, is virus sequences Minot et al. (2011). This is one of the main reasons why we need a tool that can differentiate between bacterial and viral sequences.

Current tools to know which sequences in the metagenomic data are generally using searching techniques (e.g blast). These tools assemble high throughput sequences then search again known databases with a similarity measure. One of consequences of using similarity is the detection of viruses similar to available sequenced viruses. Although, about 15% of viruses in the human gut microbiome and 10% in the ocean have similarity to the known viruses Ren et al. (2017).

Machine learning approaches are recently used in classification and clustering tasks as it provide a statistical robust technique. Machine learning systems work by learning from many examples in order to find the relation between the input and the output in classification problems and find the similarities between the input data in case of clustering problems.

Deep learning (Deep neural networks) is one of the machine learning methods that are considered as a state of the art technique for general classification problems. It shows significant improvements in several artificial intelligence tasks for example image classification, speech recognition, and natural language processing. Moreover, It shows significant results with genomic data Angermueller et al. (2016).

In this paper, we apply feature extraction methods used in natural language processing techniques that can identify viral reads robustly without any homology search and this can be used to purify viral metagenomic data from viral contamination as well. Then, we compare feature extraction with machine learning method and deep neural networks.

## 2. Related Work

Many pipelines are using BLAST Altschul et al. (1997), which is a well-known sequence alignment tool of nucleotide and protein sequences. Moreover, it finds the similarity regions between sequences with a statistical significance, which can indicate whether the target sequence belongs to the reference genome.

Other tools are using BLAST alternatives in order to have a rapid pipeline for sequence identification. DIAMOND Buchfink et al. (2014) tool is much faster and efficient than BLAST. It can align short DNA or protein sequences against protein databases. It was reported that it is 2500 times faster than BLASTX Buchfink et al. (2014).

Centrifuge Kim et al. (2016), MetaPhlAn Truong et al. (2015), and GOTTCHA Freitas et al. (2015) are used for metagenomic reads classification, as it is a modified version for BLAST similarity searches, but it is optimized for metagenomic classification problem to classify the reads with very high speed. Centrifuge uses an optimized indexing scheme based on the Burrows-Wheeler transform (BWT) and the Ferragina-Manzini (FM), which are good algorithms for data compression and pattern matching.

On the other hand, MetaPhlAn Truong et al. (2015) uses clade-specific marker genes to assign microbial reads. Clades are groups of genomes that can be as specific as species or phyla and clades. Clade-specific are the markers genes that are conserved in clade’s genomes and not has a local similarity with any sequence outside the clade. The clade marker genes are used in MetaPhlAn, are precomputed from the coding sequences, which are abundant in microbes sequences.

### 2.1. Virus-specific tools

Many available tools are using homology search with various similarity measure to known virus databases using features (e.g. genes) in order to find phages from prokaryotic genomes, such as Phage_Finder Fouts (2006), Prophinder Lima-Mendez et al. (2008), PHASTZhou et al. (2011), and PhiSpy Akhter et al. (2012). Some of them, such as PhiSpy integrates other features (e.g. unique virus k-mers, AT and GC skew, protein length and transcription strand direction).

Wang et al. 2017 have proposed VirusSeeker Zhao et al. (2017) as a robust pipeline for classification of microbial sequences. This pipeline is a BLAST-based tool that can work on eukaryotic virus discovery and virome analysis.

There are other tools, such as VIROME Wommack et al. (2012) and MetaVirRoux et al. (2011) are able to detect viral sequences in mixed metagenomic datasets. They are using similarity search against proteins. Again, using limited known reference databases is a limitation of this approach. However, These tools failed in detection of viral reads in metagenomic data because the databases are outdated, limited, and do not represent viral diversity in the environment. Moreover, It is not optimized to process a large number of contigs Roux et al. (2015a), as they depend on alignment and homology search limitations.

VirSorter Roux et al. (2015a) is able to detect phages and viral sequences. VirSorter uses similarity search to viral databases and integrates other features related to the analysis of sequence genes, such as enrichment of viral-like genes, enrichment of uncharacterized genes, and viral hallmark gene. These extracted features make the identification more accurate, but they still suffer limitations. One of the limitations, is the requirement of having at least three genes within the contig because the smallest virus genome contains three genes, because the smallest virus discovered has three genes only, so it possesses shortcoming similar to the previous techniques because of using homology strategy. Moreover, it cannot work with short fragments or contigs.

### 2.2. K-mer tools

K-mer refers to possible substrings of length k that are contained in a sequence or a string. Generally, any sequence of length *L* with *n* chars (four nucleotides in case of DNA) will have n^*k*^ total possible k-mers and the total number of k-mers is *L* − *k* + 1, where k is the length of new substrings.

K-mer is used in sequence alignment and it models the variability in the genome mutations Aggarwala and Voight (2015). Moreover, k-mer features are used in various computational biology problems, such as sequence assembly Compeau et al. (2011), improving heterologous gene expression Welch et al. (2009) and, identify species in metagenomic samples Perry and Beiko (2010); Ounit et al. (2015); Wood and Salzberg (2014). K-mers are also used in identifying biomarkers in samples related to some diseases Wang et al. (2018) and estimate the abundance of species in metagenomic data Lu et al. (2017).

In metagenomic binning algorithms, k-mer features are massively used. Binning is where separating sequencing reads into bins for each organism in order to group the contigs, reads or identify their taxonomic units Alneberg et al. (2014). For example, TETRA Teeling et al. (2004) tool bin the metagenomic reads into organisms based on their tetranucleotide where k-mer is four. Additionally, there are other various tools are used in metagenomic binning with maximum six k-mers such as TACOA Diaz et al. (2009), CompostBin Chatterji et al. (2008), PhyloPythia McHardy et al. (2007), and PCAHIER Zheng and Wu (2010).

Kraken Wood and Salzberg (2014) is an alignment based tool for annotating the metagenomic DNA sequences with taxonomic labels. It is used an exact alignment of k-mers, which help in achieving a comparable accuracy with sequence alignment tools, but it is faster than BLAST and 11 times faster than MetaPhlAn.

Clark Ounit et al. (2015) is based on reduced k-mer where *k* is more than or equals to 20, and it is five times faster than Kraken with better accuracy in classification on genus/species level. It builds an index for a target sequences first to optimize the time for k-mer alignment.

### 2.3. Machine learning tools

A machine learning tools are trained in order to assign the DNA sequences to a label (e.g. virus or bacteria). These algorithms used engineered features, such as k-mer, oligonucleotide frequency profiles, AT and GC skew, protein length, transcription strand direction, and codon usage.

Recently, VirFinder Ren et al. (2017) applies one of basic machine learning techniques, which is logistic regression. It is a statistical model that uses the k-mer features as an input. Specie’s genome varies in the number of modes in a k-mer spectrum Chor et al. (2009). K-mer similarity score can discriminate viruses within other prokaryotic genomes even the viral reads is short(500 bp). Furthermore, MARVEL Amgarten et al. (2018) tool is used for identifying bacteriophage sequences in metagenomic bins. Random forest approach is applied in MARVEL and model input is various engineered features, such as gene density, strand shifts, and fractions of significant hits to a viral protein database Grazziotin et al. (2016). It is reported that MARVEL performance is slightly better than VirFinder, but it achieves better sensitivity measure.

## 3. Materials and Methods

### 3.1. Building training and testing dataset

The data used in this study is similar to the one we used in Abdelkareem et al. (2018). It is divided into a train and test genomes randomly with 80% of total base pairs (Table 1). The available viral genomes were processed until Nov. 1^st^, 2017. whereas, prokaryotic genomes were sampled since the huge number of available prokaryotic genomes. Then, the viral genomes were converted into non-overlapping fragments of different lengths n = {100, 500, 1000, 3000}. Fragments of prokaryotic genomes were generated randomly in an approximate equal number of non-overlapping with the same lengths (Table 2). Finally, the data of both classes were balanced using a random under-sampling technique to avoid the bias to the majority class with the deep neural network. Figure 1 describe the data pipeline.

**Table 1.** The number of training and testing genomes from RefSeq.

| Genome | Train | Test | Total |
| --- | --- | --- | --- |
| Viruses | 7686 | 1870 | 9556 |
| Prokaryotes | 143241 | 35543 | 178784 |

**Table 2.** The generated viral fragments used in training and testing.

| Fragment Length (N) | Train | Test |
| --- | --- | --- |
| 100 bp | 2088863 | 527020 |
| 500 bp | 420857 | 106168 |
| 1000 bp | 212253 | 53528 |
| 3000 bp | 73163 | 18425 |

**Figure 1:**
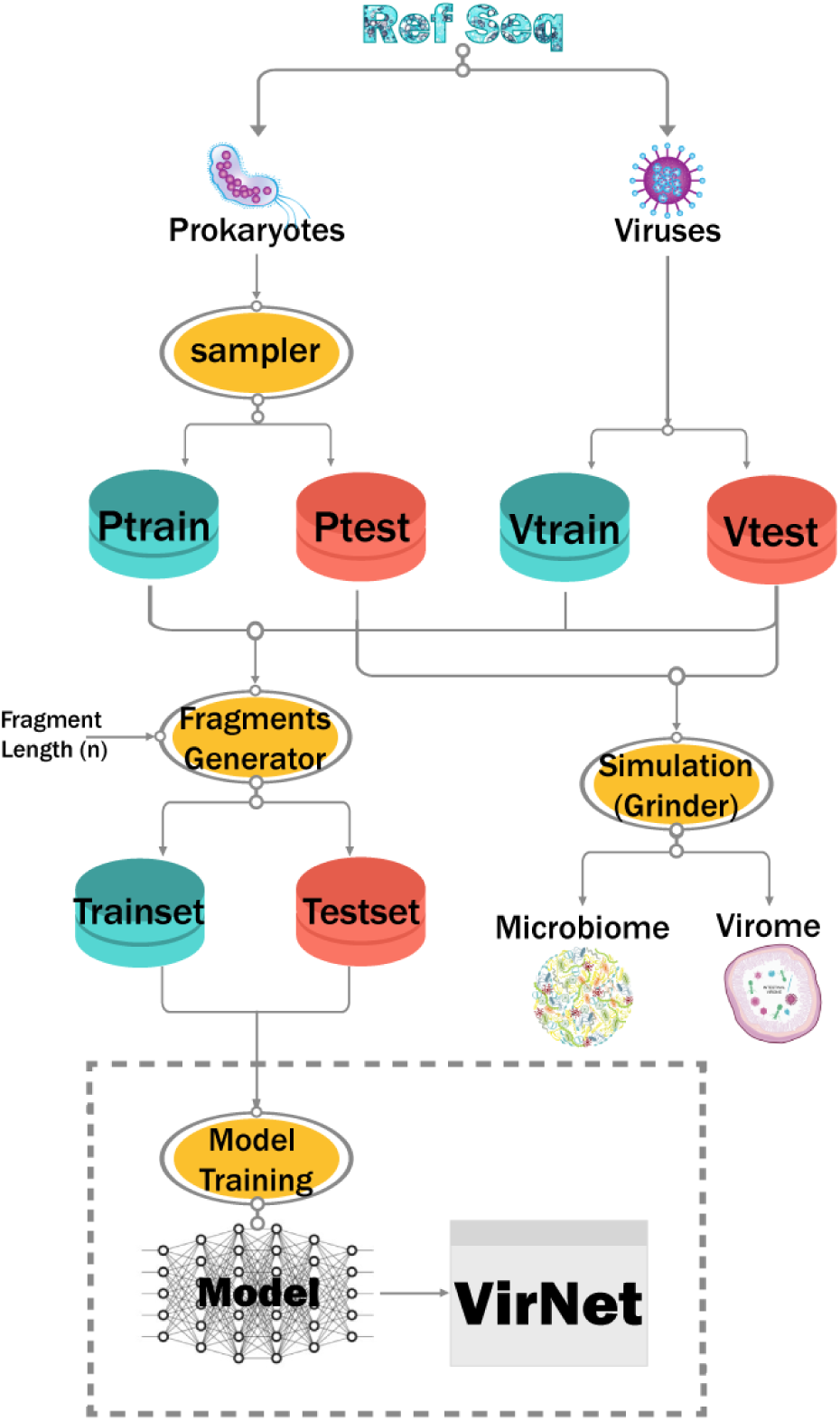
VirNet Data Pipeline.

### 3.2. Generating simulated virome and microbiome

Two metagenomic reads were generated for the purpose of the verification of the proposed tool ability to detect viral reads. Sequence of virome (with the majority of sequences from viral genomes) and microbiome with 1,000,000 reads 100 pb, were generated using Grinder Angly et al. (2012) from our reference test genomes to simulate shotgun metagenomic. The virome and microbiome data have 75% and 2% of viral reads respectively. Moreover, we used Illumina error model indicated by mutation_dist poly4 3e-3 3.3e-8 and mutation ratio 91:9 (9 indels for each 91 substitution mutations) because for Illumina indel errors occur more often than substitution errors Laehnemann et al. (2015). Table 3 shows simulated data statistics.

**Table 3.**
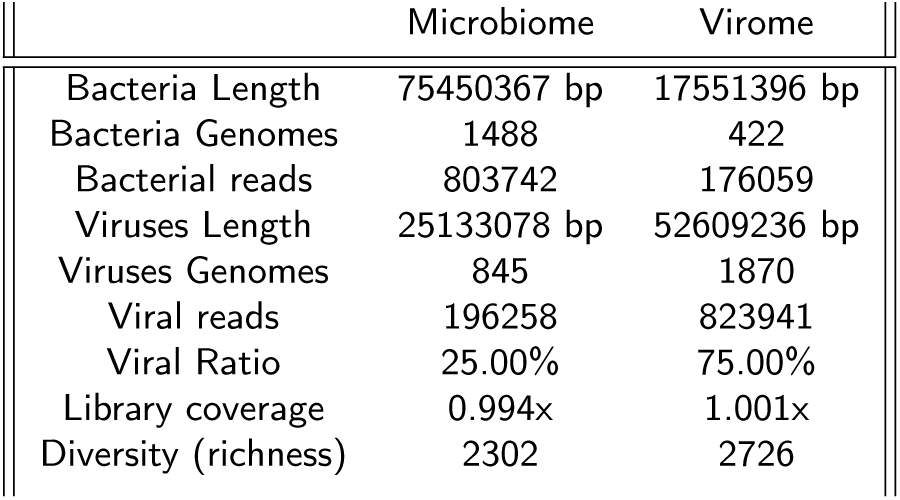
Grinder Simulated Metagenome.

### 3.3. Machine Learning Model

Viral sequence identification can be formulated as Natural Language Processing (NLP) problem Loewenstern et al. (1995). Every single nucleotide (e.g. A, C, T, G) can be considered as a word or a character in the sequence similar to the sequence of words in the natural language. Moreover, a group of nucleotide, such as consecutive three nucleotide (codon) can be treated as a word. There is an analogy between DNA sequences and text. A sequence of English words can be classified in sports category, similarly, DNA sequence of viruses can be identified as well.

Text classification is one of the essential tasks in NLP domain. There is a wide range of applications using text classification algorithms, such as topic modeling Wallach (2006), spam detection Wang (2010), language detection Jimenez-Feltström (2006), and sentiment analysis Pak and Paroubek (2010). Before building text classification models, The text has to be converted into numerical variables, which can represent the text content. Representation of this text is required based on the type of task to be applied Lewis (1992).

#### 3.3.1. Feature Extraction

The most common approach is to convert the text into bag-of-words (BOW) Joachims (1996). BOW is a vector contains the number of occurrence of each tokenized word in the text example. However, this method has some problems because it neglects the dependencies between the consecutive words. So, one of the most important feature representation for the text problems is the *n*-gram method which is considered as a contiguous sequence of *n* tokens in the input sequence Joachims (1998). This input sequence might be words, phonemes, letters or even base pairs. Also, *n*-gram models are used in many tasks because of its simplicity.

Classification task depends on the appropriate representation of documents or sequences to be classified well Quinlan (1983). There is a different representation of n-gram features, such as Term Frequency (TF) and Term frequency-Inverse document frequency (TFIDF) Ramos et al. (2003).

#### 3.3.2. Logistic Regression

Supervised learning is one of the machine learning tasks which try to find the relation or the function which map the input example with the output. There is a wide range of algorithms such as logistic regression, decision trees, random forest, support vector machines (SVM), and neural networks. Logistic regression is a statistical technique used in machine learning for classification problems when the target value is categorical and it used in many biological and economical applications.

The learning algorithm in logistic regression is trying to find the values of the optimal coefficients *b* using maximum-likelihood estimation or stochastic average gradient descent (SAG). SAG method is much faster for large datasets. Most machine learning algorithms for classification problems evaluated by accuracy. However, accuracy is not sufficient as your testing data might not being balanced.

In this section, we will demonstrate several machine learning evaluation metrics used in experiments. Let *T P* is the number of predictions where the predicted samples are positive and the actual samples are positive. *T N* is the number of predictions where the predicted samples are negative and the actual samples are negative too., *P*, and *N* are the total number of positive and negative samples respectively.

- Accuracy (ACC): It is the ratio of correct predictions to the total number of the examples. For binary classification

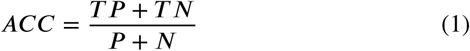

as we mentioned accuracy fails to evaluate the classification model when the number of samples of each class is not approximately equal.
- Precision (PPV): It is also known as positive predictive value and it is the ratio that indicates how relevant our prediction to the whole retrieved data and this metric shows how many items are relevant.

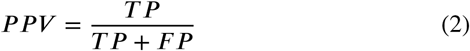

where *F P* is the number of predictions where the predicted samples are positive and the actual samples are negative.
- Recall (TPR): It is also known as a true positive rate (sensitivity) and it is the ratio that indicates how relevant our prediction to the whole retrieved data and this metric shows how many relevant items are selected.

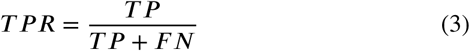

where *F N* is the number of predictions where the predicted samples are negative and the actual samples are positive.
- F-Score: is the harmonic mean of precision and recall.

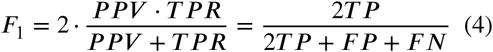 It measures the effectiveness of retrieval and relevance and the greater the F1 score, the better is performance.
- Area under Curve (AUC): It is the area under the receiver operating characteristic curve (ROC) which is the plotting of true positive ratio (TPR) and false positive ratio (FPR) with different threshold points. It shows the ability of the classifier to predict a randomly chosen positive examples higher than negative ones. The value of AUC is in the range [0,1].

TF and TFIDF are commonly used in information retrieval problems. We tried the same approach using common machine learning techniques with different n-grams settings. Table 4 and Figure 2 shows the evaluation results using n-grams equals to 5. It shows also that logistic regression (LR) outperforms other classifiers such as Xgboost, AdaBoost, Random forest (RF) and decision trees (DT).

**Table 4.**
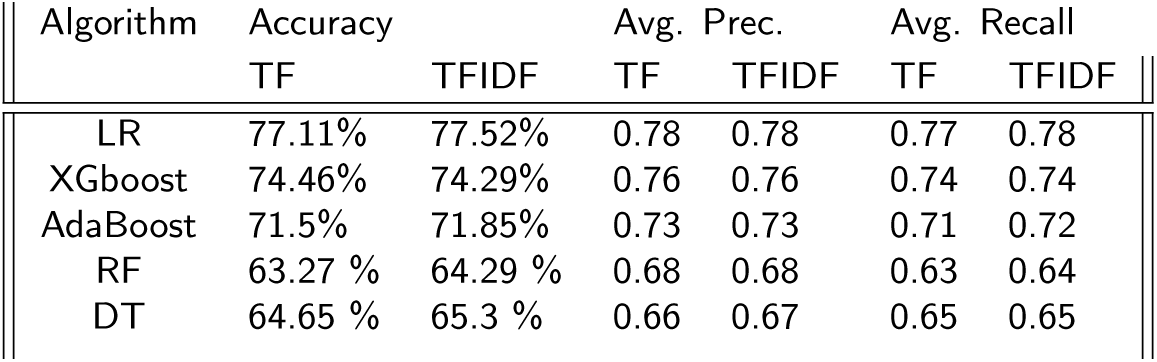
Results of different algorithms using TF and TFIDF featurizers on 5 n-grams.

**Figure 2:**
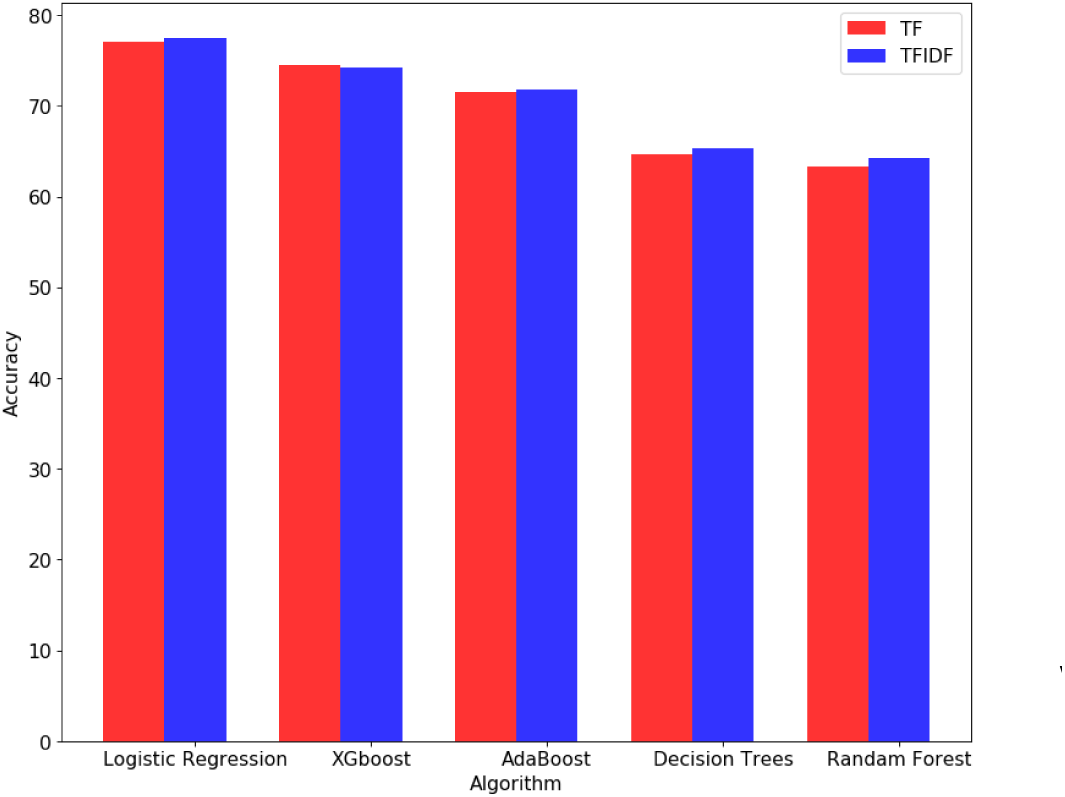
Comparison between different algorithms on accuracy using TF and TFIDF featurizers on n-grams=5 only

Moreover, we tried different n-grams in the range [1, 20] in order to find the optimal n-grams settings using logistic regression only. Figures 3 and 4 show that with n-grams=11 and TFIDF featurizer we can get 88% accuracy on the test data of 500 bp.

**Figure 3:**
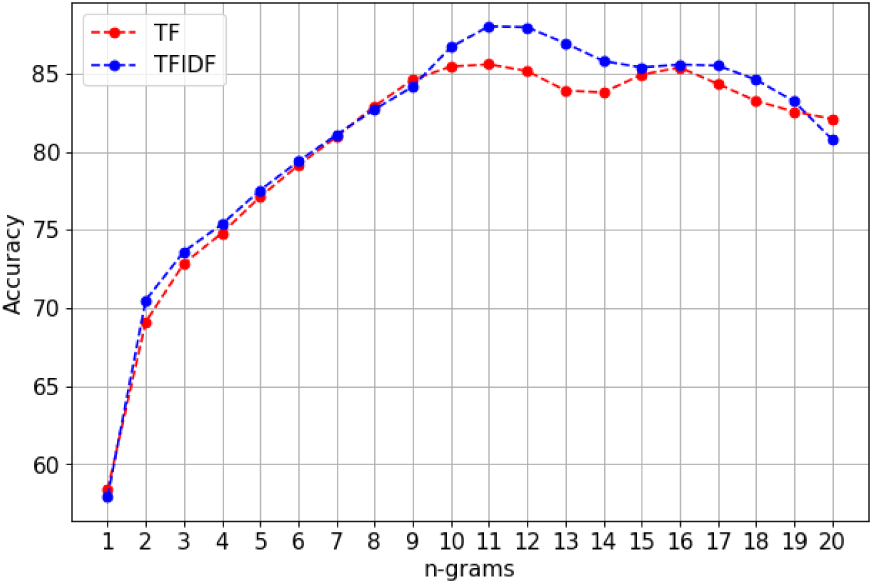
Comparison between effect of increasing n-grams on accuracy using TF and TFIDF featurizer.

**Figure 4:**
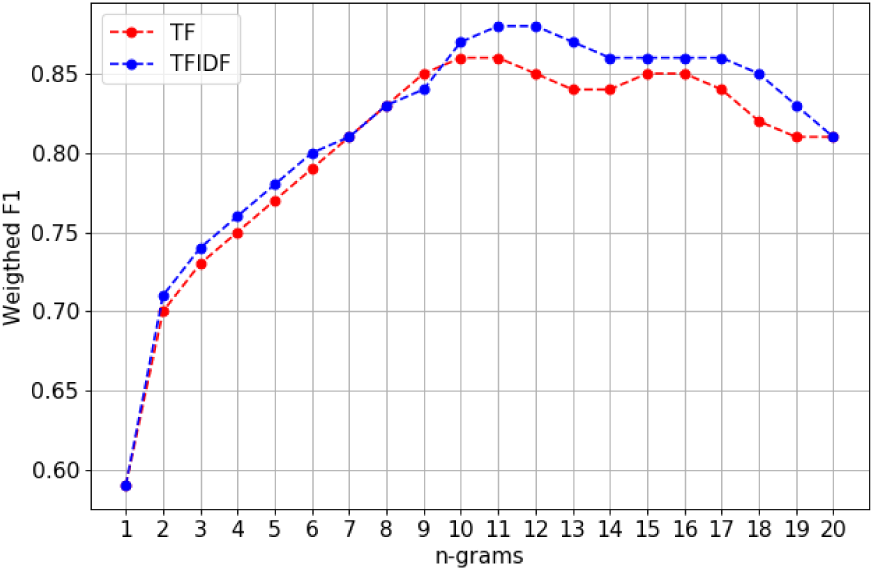
Comparison between effect of increasing n-grams on weighted F1 score using TF and TFIDF featurizer.

Regarding the speed of training and prediction, changing the n-grams could affect the training speed dramatically but we find that TFIDF has a lower training speed than TF. Figures 5 and 6 show the effect of n-grams on training time and prediction time respectively. We can conclude that there is a slight difference between TF and TFIDF in prediction time too.

**Figure 5:**
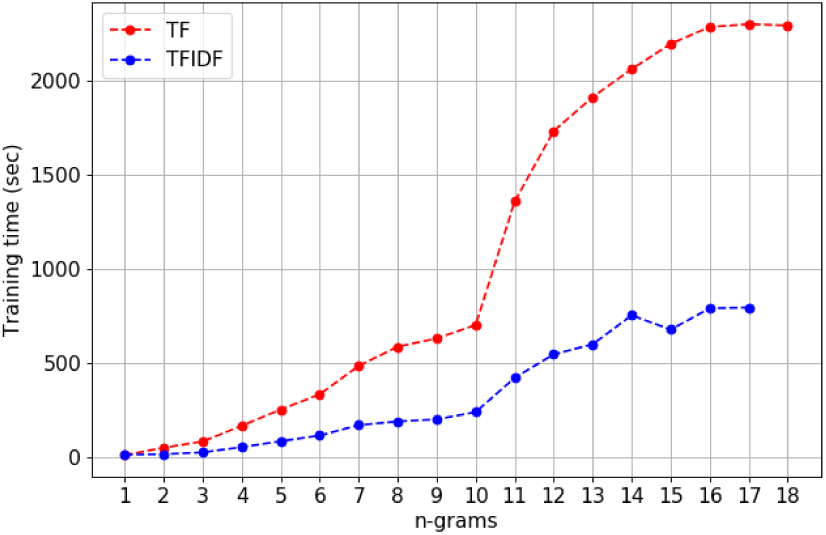
Comparison between effect of increasing n-grams on training time using TF and TFIDF featurizer.

**Figure 6:**
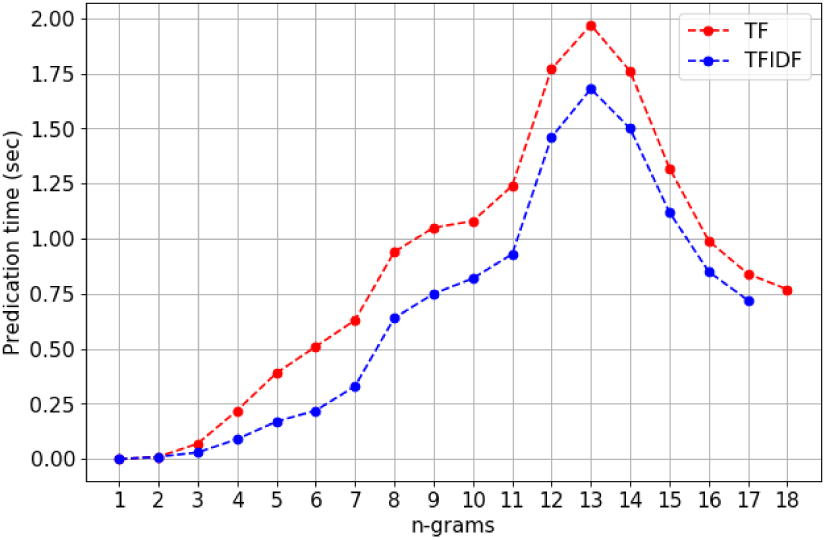
Comparison between effect of increasing n-grams on prediction time using TF and TFIDF featurizer.

#### 3.3.3. Feature Importance

Feature importance or selection is a necessary step in machine learning. Using these methods effectively could decrease the training time and improve accuracy by removing the redundant and unimportant features Jing et al. (2002). A feature is considered not important if changing its value makes the model error unchanged. Logistic regression interpretation is very simple as it based on the weighted sum of its features. There are 4,268,111 features of 11 n-grams and this is less than all combinations of our five nucleotides (A, C, T, G, and N) which equals to 5^11^ (around 8.7% of all combinations) and approximately all combinations for four nucleotides without N (noisy).

In order to find the most important features, all coefficients were sorted. The relative feature importance were checked for the best experimental setting from the previous experiments (11 n-grams, TFIDF featurizer, and Logistic regression classifier) in order to get the most important features in classification. Figures 7 and 8 show the top and bottom features respectively. These figures present different fragments, which can be investigated and checked in further studies. These fragments might be very important in order to identify viruses based on them. In case, they are essential for viral identification, practical and biological studies should be applied to prove that in lab work.

**Figure 7:**
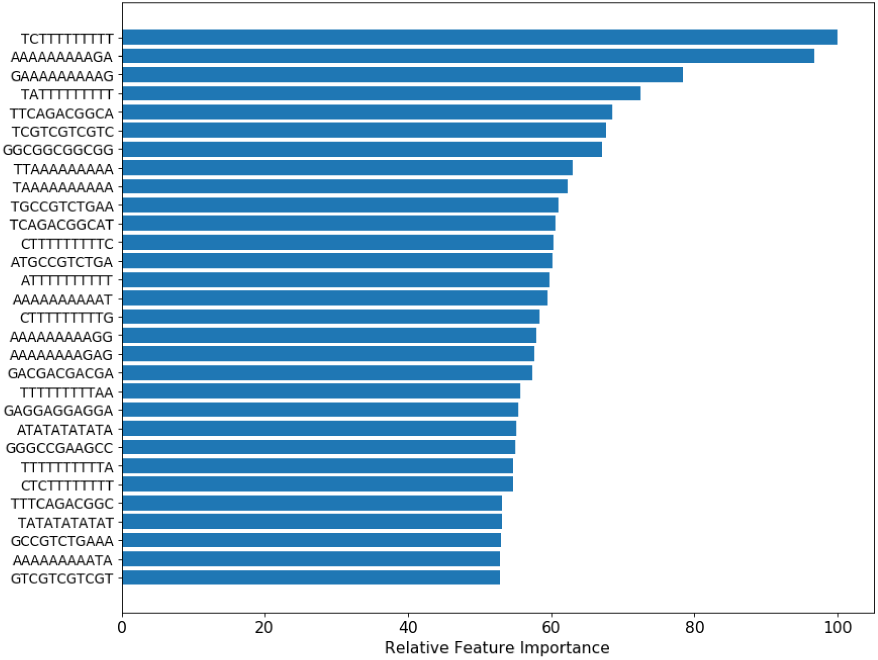
Top 30 features for TFIDF using logistic regression.

**Figure 8:**
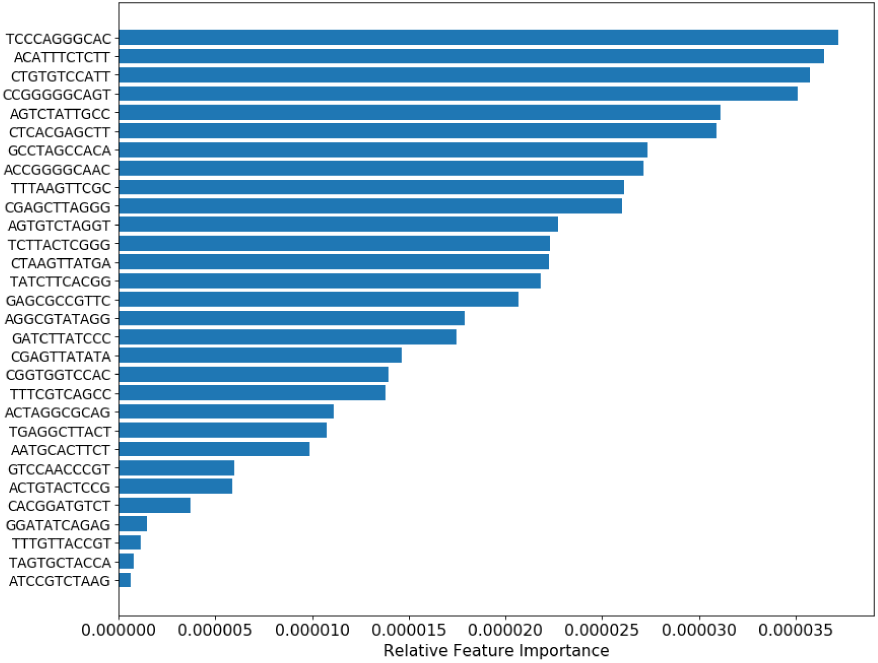
Bottom 30 features for TFIDF using logistic regression.

### 3.4. Deep Learning Model

Sequence modeling problems such as DNA sequences can be modeled using Recurrent neural networks (RNN), long short-term memory (LSTM) Hochreiter and Schmidhuber (1997) and gated recurrent neural networks (GRU) Chung et al. (2014) because they are deep neural networks architecture suitable for modelling complex relationships. Embedding layer maps discrete input words to dense vectors for computational efficiency before feeding this sequence to LSTM/GRU Layers.

LSTM encoder has three gates to protect and control the cell state, the input gate denoted **i** which defines how much of the newly computed state, forget gate denoted **f** that decides what information is to be kept and what is to be thrown away, the output of the update gate denoted **U** that’s used to update the cell state and the output of the LSTM cell **o**. **W** is the recurrent connection at the previous hidden layer and current hidden layer and **C** is the internal memory of the unit as shown in the following equations

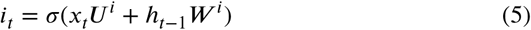

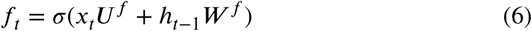

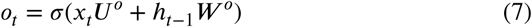

GRU encoder is same as LSTM except it has only 2 gates, Reset gate denoted **r** that determines how to combine the new input with the previously saved input state and the update gate denoted **z** that defines the amount of information to keep around, as defined in the following equations

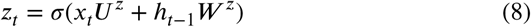

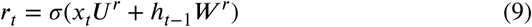

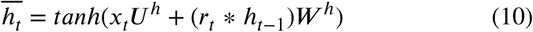

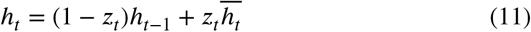

The attentional network could be applied to the current problem as learn how to model non-linear relationships and extract appropriate features from the raw data because the network will attend to previous DNA nucleotide within the same input sequence. The attentional neural model used in the proposed architecture, was trained with the DNA nucleotide bases with fragments of various lengths. The model output predictions are in binary output format whether the fragment predicted as a viral or not.

The best experimented neural network model (Figure 10b) consists of an input embedding layer of size 128 mapping input DNA nucleotide tokens into an embedding space, that is fed to an LSTM layer. The forward sequence 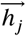 is then averaged together to create an attentional vector representing token context within the same fragment. A dropout layer was added after the attentional layer to avoid overfitting over the input data. This model is available on github as an open-source project^4^.

**Figure 9:**
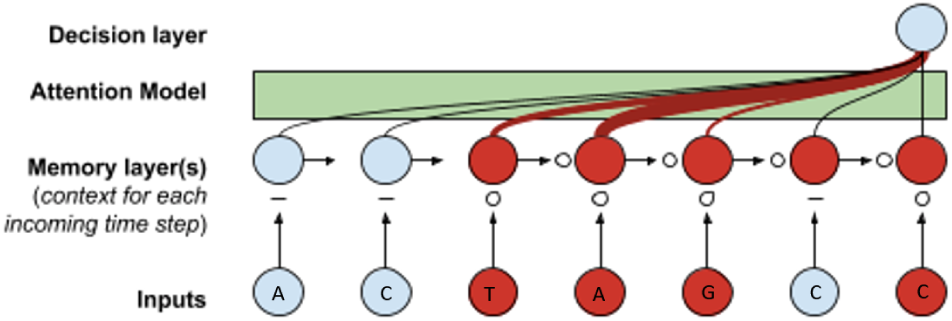
Attention mechanism with DNA sequences.

**Figure 10:**
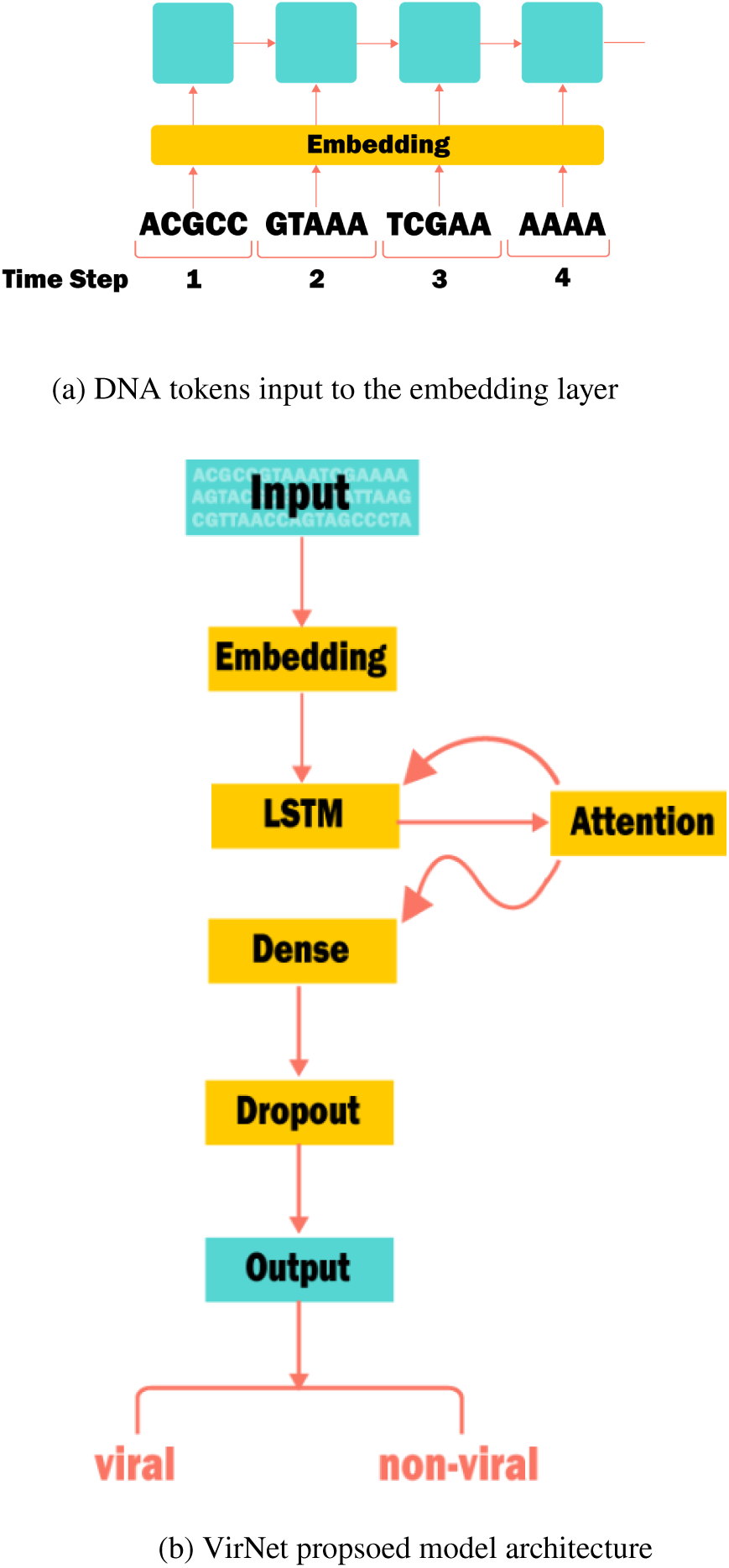
VirNet model.

LSTM layer was performing better as shown in Table 5 than the GRU cell. GRU encoder having fewer gates than LSTM model make it faster and easier to converge but depending on the size and the format of the input data. LSTM with more gates would be slower but will outperform GRU encoder type. The input sequence is divided into tokens with size 5 bp each (5-grams) and treated as a single word (Figure 10a), which mapped as a point in the embedding space. Moreover, Adam optimizer was used during the training process to maximize the conditional probability of tokens found together.

**Table 5.** Hyperparamters optimization results.

| #Layers | Embedding | Ngram | ROC-AUC | Accuracy |
| --- | --- | --- | --- | --- |
| 1 | 32 | 3 | 0.8 | 73.66 |
| 1 | 32 | 5 | 0.83 | 76.42 |
| 1 | 32 | 7 | 0.79 | 72.11 |
| 1 | 64 | 3 | 0.83 | 76.1 |
| 1 | 64 | 5 | 0.83 | 75.98 |
| 1 | 64 | 7 | 0.77 | 69.96 |
| 1 | 128 | 3 | 0.83 | 75.76 |
| 1 | 128 | 5 | 0.85 | 77.41 |
| 1 | 128 | 7 | 0.78 | 70.93 |
| 2 | 32 | 3 | 0.8 | 73.25 |
| 2 | 32 | 5 | 0.83 | 76.49 |
| 2 | 32 | 7 | 0.79 | 72.82 |
| 2 | 64 | 3 | 0.81 | 73.96 |
| 2 | 64 | 5 | 0.84 | 76.46 |
| 2 | 64 | 7 | 0.78 | 72.53 |
| 2 | 128 | 3 | 0.83 | 76.15 |
| 2 | 128 | 5 | 0.85 | 77.9 |
| 2 | 128 | 7 | 0.78 | 70.63 |

In this model, an early stopping mechanism was used as a form of regularization to avoid over-fitting over the data while making more epochs over the data. The early stopping mechanism was used with patience of three non-improving consecutive epochs; the neural model will stop training while saving the latest improving checkpoint over the validation set defined.

### 3.5. Attention mechanism

Sequence networks contain encoder layer to decode the variable inputs into a fixed-length vector. Then, it outputs a translation from the encoded vector using the decoder layer. The neural network is not able to model the relation between the input and the output and cannot compress it into a single fixed-length vector. To solve this issue, the attention mechanism was introduced in machine translation domain Bahdanau et al. (2014) to represent long sentences. It is frequently used in sequence to sequence models in machine translation but Yang et al. (2016) is used in document classification problems as well. The attention mechanism makes the neural network to focus on relevant parts in the input more than the irrelevant parts. It allows the model to learn the generation of the context vector as well.

#### 3.5.1. Context Vector

The context vector **c**_**i**_, while **a** is the alignment model, which is a feedforward neural network that is trained with all the other components of the proposed system. The alignment model scores **e** how well each encoded input **h** matches the current output of the decoder **s**. The alignment scores are normalized using a softmax function. The context vector is a weighted sum of the annotations **hj** and normalized alignment scores.

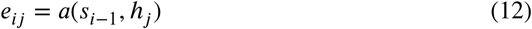

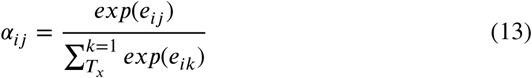

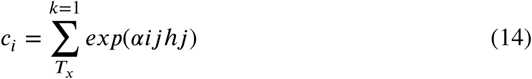

#### 3.5.2. Attention types

There are two types of attention that can define the mode of attention in the model, such as soft attention and hard attention.

1. In soft attention, the attention is able to access the entire input Bahdanau et al. (2014). In this type, the alignment weights are learned from all parts in the input so the model will be differentiable. On the other hand, this approach is a very expensive approach and it appears in huge training datasets and it shows a slower time in model inference.
2. In hard attention, it selects only one part in the input and attends to at each step of the output generation as presented in Luong et al. (2015), so the model is faster in inference time due to fewer calculations but it requires variance reduction in the training phase.

Luong, et al., 2015 Luong et al. (2015) shows another two types of global and local attention to reduce the gap between hard and soft mechanisms. The global attention is similar to the soft attention, while the local make the model differentiable because the local attention focuses on a selected small part in the input source.

#### 3.5.3. Attention in the proposed tool

DNA sequences can be considered as a sequence of words. The similarity between each two sequences is known by sequences alignment techniques. There are a similarity between the alignment in machine translation Och and Ney (2004) and DNA sequences Altschul et al. (1990). Two sentences in different languages have similar meanings even they have not the same words. In order of that, the virus DNA sequences can be formulated in the same way.

Attention mechanism is widely used in machine translation field Klein et al. (2017); Luong et al. (2015); Vaswani et al. (2017); Firat et al. (2016). Similarly, attention mechanism can be used with viral identification task to allow the model finding important features while training the recurrent neural network. In biology, sequence motif is a sequence of nucleotide that is has a biological significance in the DNA sequence Bailey et al. (2009). There are many unknown input sequences can affect the model performance. Some of DNA sequence have known meaning such as stop codons. Therefore, attention encoder focuses relevant DNA nucleotides more than the irrelevant DNA nucleotides as shown in Figure 9, relevant parts are in red.

The proposed model is implemented as an attentional encoder network (Figure 10a). An input sequence **x** = (**x**_**l**_, …, **x**_**m**_) and calculates a forward sequence of hidden states 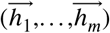. The hidden states 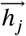 are averaged to obtain the attention vector **h**_**j**_ representing the context vector from the input sequence.

### 3.6. Hyperparameters optimization

Model parameters affect the performance of the deep learning model, and they control the behavior of the training algorithm as well. Several experiments are applied to the model using brute force search with different combinations. These experiments were on 20% of our training data with a read length 500 bp in order to run more experiments in a shorter time. The best parameters of our model including the number of recurrent layers, the embedding size for each layer and the input sub-words (ngram), are experimented and reported in (Table 5). It shows that the best setup is the model with two layers, 128 embedding size, and five ngram as an input token.

We ran other experiments with the same parameters to check the ability of our model with other configuration, so we changed different parameters separately such as embedding size to 256 neurons, the number of layers to 3 and the recurrent cell type to GRU instead of LSTM. We found a slightly less accuracy by 0.01% and ROC-AUC scores.

## 4. Results

### 4.1. Results for generated fragments

VirNet was experimented fragments with various lengths n= {100, 500, 1000, 3000} bp. These fragments were generated from our testing set of viruses and prokaryotes RefSeq genomes. VirNet outperformed VirFinder on all fragments lengths. The results are showed in Figure 11. It presents the ability of VirNet model to predict viral reads in mixed metagenomic data. The accuracy of VirNet was **82.82%** on 500 bp, whereas VirFinder was **75.61%**.

**Figure 11:**
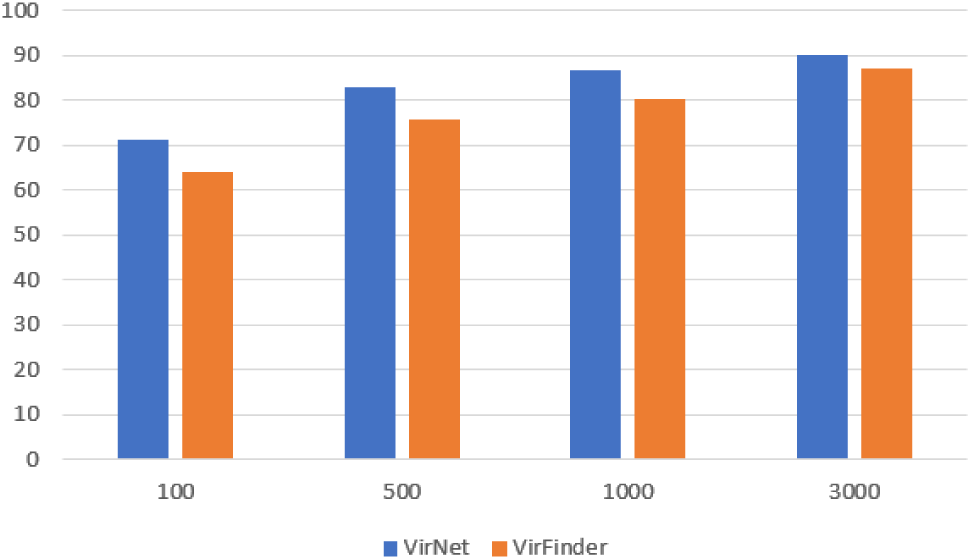
VirNet vs VirFinder Accuracy.

Furthermore, VirNet was able to predict the short fragments with length 100 bp by **71.29%**, while VirFinder (VF) is not able to identify viral reads less than 500 bp because it was trained by 500 bp as a shortest fragment length. It is observed from Figure 11 as well, the sensitivity of our model to the length of fragment. The accuracy was dropped by decreasing the fragment’s length.

The average precision and recall are similar because of balancing the both classes data. Table 6 presents the accuracy, average precision, and average recall of both tools. It also shows, VirNet of average precision and recall equal to **0.83** on length of fragment 500 bp, while **0.76** in VirFinder.

Receiver Operating Characteristic (ROC) curves were plotted for both VirFinder and VirNet prediction results on the same testing set (Figures 12a and 12b). ROC curve is a plot of the true positive rate against the false positive rate for the different possible thresholds. Moreover, curve is in the upper left corner, which means higher overall accuracy of both classes (viruses and prokaryotes). It shows an increase in area under the curve (AUC) by increasing the length of fragments. It is reported AUC of **0.69, 0.82, 0.85**, and **0.90** on fragment length of 100, 500, 1000, and 5000 respectively using VirFinder (Figure 12a), whereas VirNet is **0.79, 0.88, 0.93**, and **0.92** (Figure 12b).

**Table 6.**
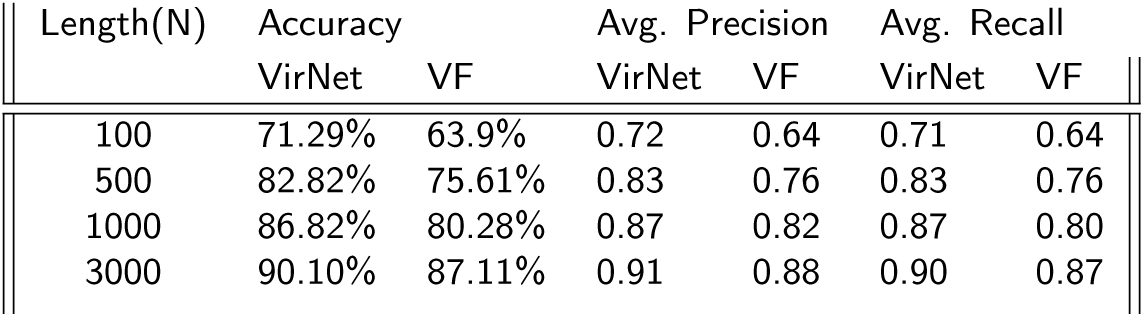
Comparison on fragments test-set.

| Length(N) | Accuracy |  | Avg. Precision |  | Avg. Recall |  |
| --- | --- | --- | --- | --- | --- | --- |
|  | VirNet | VF | VirNet | VF | VirNet | VF |
| 100 | 71.29% | 63.9% | 0.72 | 0.64 | 0.71 | 0.64 |
| 500 | 82.82% | 75.61% | 0.83 | 0.76 | 0.83 | 0.76 |
| 1000 | 86.82% | 80.28% | 0.87 | 0.82 | 0.87 | 0.80 |
| 3000 | 90.10% | 87.11% | 0.91 | 0.88 | 0.90 | 0.87 |

**Figure 12:**
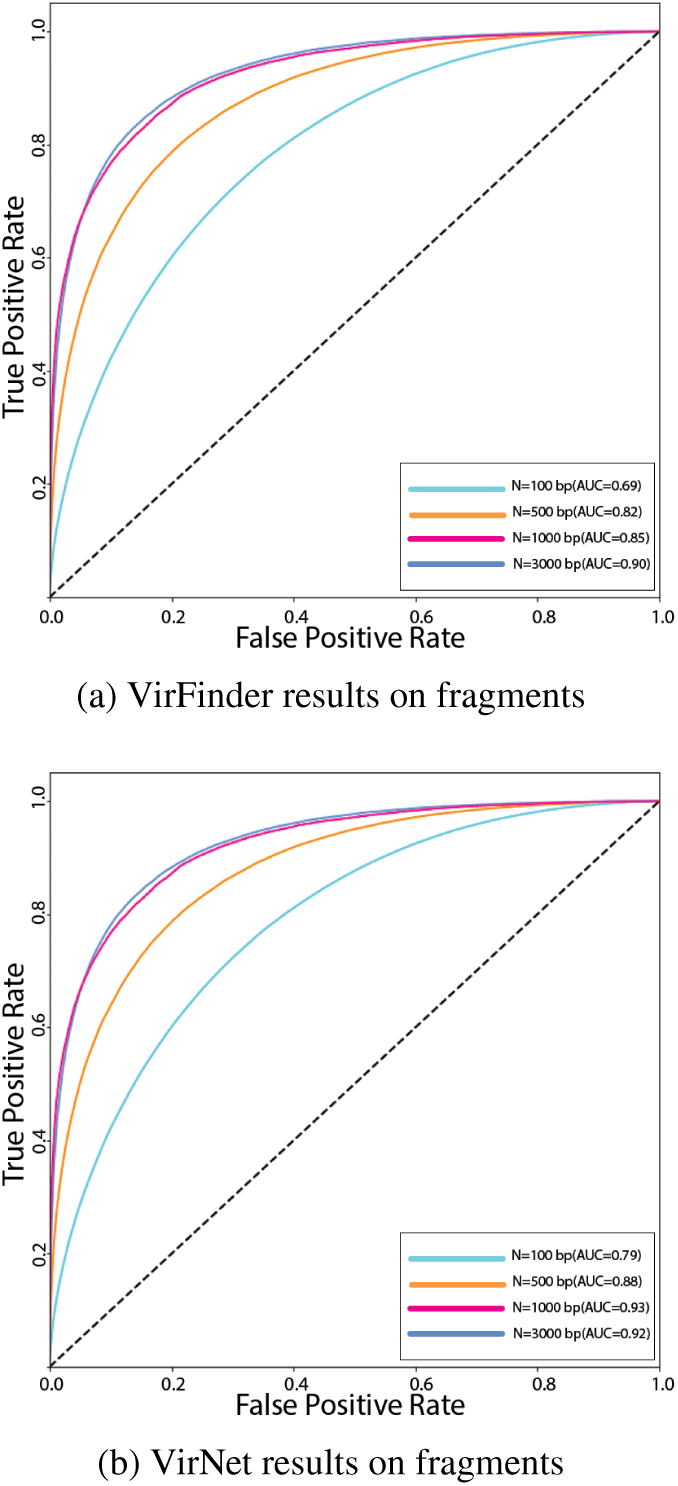
ROC-AUC curves for VirFinder on fragments.

### 4.2. Results for simulated metagenomes

VirNet outperformed VirFinder on short reads simulated metagenomes. VirNet accuracy was **71.3%** on the virome data and **72.14%** on the microbiome, On the other hand, VirFinder is **62.77%** and **64.49%** on the same data respectively.

Table 7 presents the comparison between VirNet and VirFinder in terms in accuracy, average precision and average recall. These results make sense as the accuracy is around 70%, which is similar to reported results on the testing dataset for fragments of length 100 bp, because virome and microbiome data are 100 bp as well. Furthermore, it presents VirNet of average precision and recall equal to **0.71** and **0.73** in microbiome and virome data respectively, while **0.63** and **0.65** using VirFinder, likewise the results on testing fragment dataset with length 100bp.

**Table 7.** Simulated Metagenomes Results.

| Metagenome | Accuracy |  | Avg. Precision |  | Avg. Recall |  |
| --- | --- | --- | --- | --- | --- | --- |
|  | Net | VF | Net | VF | Net | VF. |
| Virome | 71.3% | 62.77% | 0.71 | 0.63 | 0.72 | 0.62 |
| Microbiome | 72.14% | 64.49% | 0.73 | 0.65 | 0.73 | 0.64 |

Similarly, ROC curves were plotted for both VirFinder and VirNet prediction results on the simulated metagenomes testing set (Figures 13b and 13a). The curve shows high overall accuracy, sensitivity, and specificity of prediction of both classes (viruses and prokaryotes). It is reported AUC of **0.70** and **0.68** on microbiome and virome data respectively using VirFinder (Figure 13a), whereas VirNet is **0.75** and **0.74** (Figure 13b).

**Figure 13:**
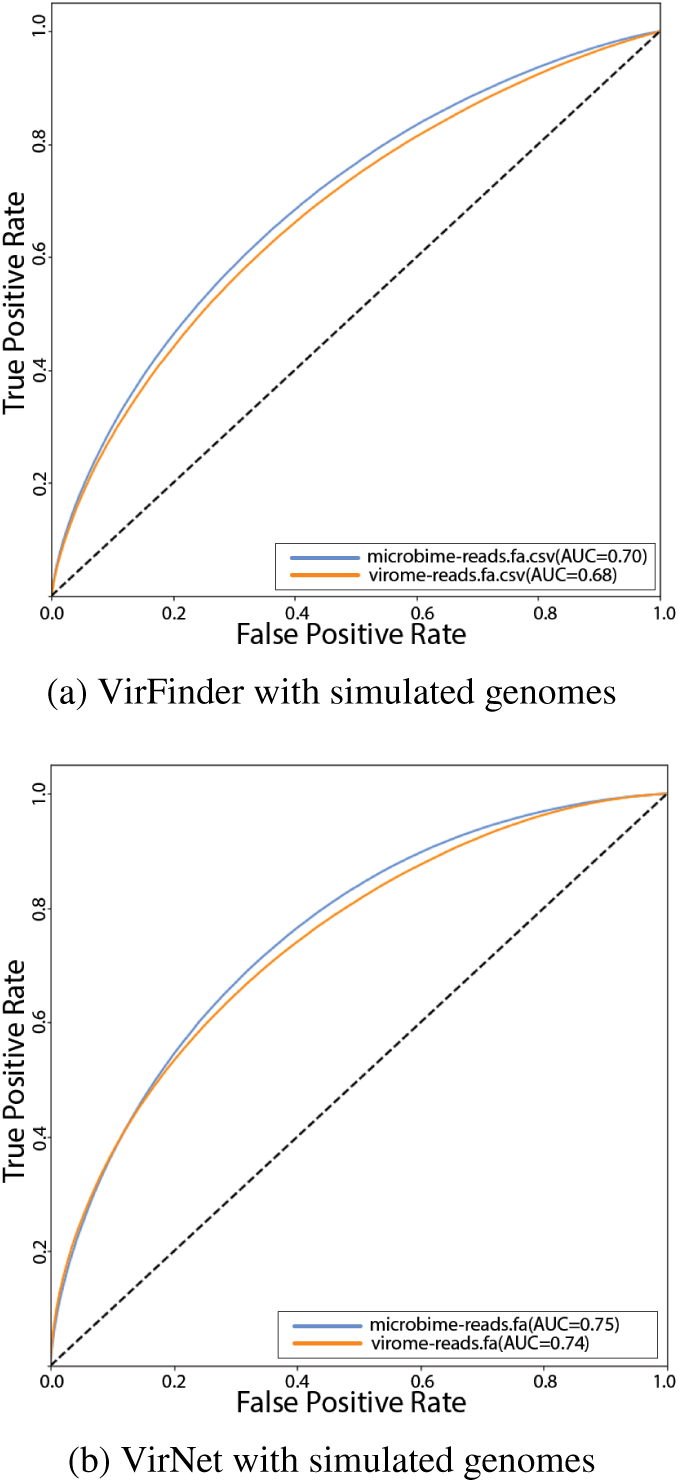
ROC-AUC curves on simulated metagenomes.

### 4.3. Results for real data

The tool was experimented on two real metagenomic data generated by two different machines (Roche 454 and Illumina). Table 8 reports an approximate equal number of viral reads in Lake Michigan virome data in both pair ended files. These results prove the ability of VirNet tool to work on pair ended reads, which will increase the sensitivity of the prediction by neglecting the conflict between pair-ended reads.

**Table 8.** VirNet results on real data.

|  | SRR648314 | SRR995836/1 | SRR995836/2 |
| --- | --- | --- | --- |
| Viral | 39,447 | 579,697 | 590,869 |
| Non Viral | 20,866 | 1,055,975 | 1,044,803 |
| Total | 60,313 | 1,635,672 | 1,635,672 |

## 5. Discussion

In the proposed tool, The network will learn how to extract suitable features from the raw data without any engineered processing to sequences (e.g. K-mer). It reveal non-linear relationships between the raw input data and the model can be used to different kind of genomes. This model is more accurate than current available tools based on our experiments. The short viral sequences were identified because LSTM learns from the dependencies between the input sequence. For the trained deep learning model, using a sliding window over the input DNA sequence might improve the proposed model, the only drawback of this technique is the slow training and inference time of input sequences. Also, using an adaptive learning rate decaying over time steps during the training might improve the model performance, but it needs more tuning over the input data.

Additionally, TFIDF can be used instead of k-mer in reads identification as they indicate a unique for each classification class. Finally, according to these results, using shallow model with TFIDF feature can give us better results than deep attention model. It is not clear that the deep sequence models are the best option for DNA sequence classification problems. Although deep neural networks have in theory much higher representational. This results is similar to the result published in Mikolov et. al 2016 Joulin et al. (2016) on a similar problem (text classification) and same model architecture.

## 6. Conclusion

We could successfully implement neural networks in viral read identification with better accuracy that current methods, which could improve the future analysis of metagenomics data and might help identify novel viruses. Although, various methods using homology and statistical techniques are able to identify viral sequences. These methods encounter many limitations because of the limited genomic databases and viral genome diversity.

In this work, It reveals the importance of using the text feature extraction pipeline to transform DNA base pairs into a set of characters with a term frequency and inverse document frequency featurization technique. Furthermore, an attention deep neural network model was introduced for identifying viral reads inside the metagenomic data. This method is indispensable to purify mixed metagenomic data from viral contamination.

The proposed network succeeded in short fragments identification. The deep neural network approach was validated with an accuracy of more than **83%**. Moreover, TFIDF was experimented to consider as a better signature for organisms instead of TF or (k-mer). By converting DNA sequence as char ngrams in form of TF or TFIDF features in a deep feed-forward neural network, it outperformed the attention model with an accuracy more than **89%**. According to these results, the proposed model would help to understand viruses in various microbial communities and discovering new species of viruses.

## Footnotes

4 https://github.com/alyosama/virnet

